# Factor VII activating protease (FSAP) inhibits the outcome of ischemic stroke in mouse models

**DOI:** 10.1101/2022.01.12.476006

**Authors:** Jeong Yeon Kim, Dipankar Manna, Trygve B. Leergaard, Sandip M. Kanse

## Abstract

Factor VII activating protease (FSAP) is a circulating serine protease, and individuals with the Marburg I (MI) single nucleotide polymorphism (SNP), which results in an inactive enzyme, have an increased risk of stroke. The outcome of ischemic stroke is more marked in FSAP-deficient mice compared to their wild-type (WT) counterparts. Plasma FSAP levels are raised in patients as well as mice after stroke. *In vitro*, FSAP promotes fibrinolysis by cleavage of fibrinogen, activates protease-activated receptors and decreases the cellular cytotoxicity of histones. Since these are desirable properties in stroke treatment, we tested the effect of recombinant serine protease domain of FSAP (FSAP-SPD) on ischemic stroke in mice. A combination of tissue plasminogen activator (tPA) and FSAP-SPD enhanced clot lysis, improved microvascular perfusion and neurological outcome and reduced infarct volumes in a mouse model of thromboembolic stroke. In the tail bleeding model FSAP-SPD treatment provoked a faster clotting time indicating that it has a pro-coagulant effect that is described before. FSAP-SPD improved stroke outcome and diminished the negative effects of co-treatment with tPA in the transient middle cerebral artery occlusion model. The inactive MI-isoform of FSAP did not have any effects in either model. In mice with FSAP deficiency there were minor differences in the outcomes of stroke but the treatment with FSAP-SPD was equally effective. Thus, FSAP represents a promising novel therapeutic strategy in the treatment of ischemic stroke that requires further evaluation.

## Introduction

An abrupt loss of brain blood supply due to arterial thrombosis is the major cause of ischemic stroke and it afflicts a large number of people worldwide^1^. Along with the surgical removal of clots from larger arteries, the only approved pharmacological therapy is tissue plasminogen activator (tPA)^2^. tPA treatment is effective in the 0-4.5h time frame, after which the adverse effects outweigh the positive attributes^2^. Moreover, it is efficacious in only 30% of the patients^3^, and its use is associated with hemorrhagic transformation in about 5-10% of cases^4^. Thrombectomy is effective over a longer time range, it can be combined with tPA and has a better cost-benefit ratio^5^. However, its widespread implementation will require a considerable investment of resources. Hence, there is an urgent need to expand the time-window of treatment, improve the effectiveness of tPA and reduce its serious side-effects.

Factor VII activating protease (FSAP) is a circulating serine protease, and individuals with single nucleotide polymorphisms (SNPs), in or near the FSAP-gene locus, have an increased risk of stroke^6,7^. Of these SNP’s, the Marburg-I (MI) SNP, results in a protein with low proteolytic activity^8^. Circulating FSAP levels are increased in patients with ischemic stroke^9,10^. It has also been observed that increased plasma FSAP levels correlate with poor neurological outcome^9^. Mice have elevated levels of FSAP in the brain and plasma after stroke^9^ and FSAP-deficient mice subjected to thromboembolic stroke exhibit a larger infarct volume and increased behavior deficit than wild type (WT) mice^11^.

Although the activation of FVII has been the name-giving property of this protein^8^, a number of other activities have been ascribed to it in relation to thrombosis and hemostasis. It can activate pro-urokinase (uPA)^8^, cleave TFPI and decrease its activity^12^, and cleave fibrinogen and speed up lysis of clots^13^. In light of these antagonistic effects on coagulation it is difficult to ascertain its net effect *in vivo*. The MI-SNP has not shown to be decisively linked to thrombosis^14^ and FSAP-deficient mice exhibit delayed thrombosis as well as hemostasis^15^. FSAP is a pleiotropic protease with a broad substrate specificity^16^. Amongst its many effects, it stimulates protease activated receptors (PARs) and increases vascular permeability in the lungs^17^ as well as inhibiting apoptosis^18^. FSAP is exceptionally effective in blocking the toxicity of histones^19^ and is more potent in this activity than activated protein C (APC)^20,21^. Histones are released during necrosis, apoptosis or in neutrophil extracellular trap (NETosis) formation^22^ and are potent activators of the zymogen form of FSAP^19^. Thus, histones trigger a self-destructing cycle that leads to their own neutralization via FSAP.

Thus, converging lines of evidence from human genetic investigations, mouse models and *in vitro* studies^6,11,13,18,23^ indicate that FSAP may be a candidate for the therapy of ischemic stroke. A recent study shows that high molecular weight-hyaluronic acid, which is a pleiotropic factor with multiple functions^24^, downregulates endogenous FSAP expression and improves the outcome of ischemic stroke in mice^9^. This indirectly implicates FSAP in the outcome of stroke in mouse models. We have recently expressed the recombinant serine protease domain (SPD) of human FSAP and confirmed that it recapitulates all, the tested, activities of the endogenous full-length protein^25^. Here, we report on the efficacy of FSAP-SPD treatment in mouse models of ischemic stroke.

## Results and Discussion

Before testing FSAP-SPD in vivo, we performed a pilot pharmacokinetic study to gain information about its half-life *in vivo*. Wild type (WT)-FSAP-SPD was measured in plasma by Western blotting, human FSAP-specific enzyme-linked immunosorbent assay (ELISA) as well as by using a fluorogenic substrate specific for the enzymatic activity of FSAP^26^. ELISA and Western blotting indicated a shorter half-life (2-3 min) compared to the activity based assay (4-5 min) in mice (Suppl. Fig, 1a-d). Similarly short half-life times have also been reported for tPA^27^, and are probably due to complex formation of proteases with inhibitors in the blood and their subsequent cellular uptake. Based on these results we elected a 30-min infusion protocol for the application of WT-SPD.

We first used a model of thromboembolic stroke (TES) where thrombosis was induced in the carotid artery with a topical application of FeCl3 which causes oxidative damage to the vessel wall^28^. A primary thrombus blocks the artery while microthrombi embolize to the brain microvasculature^29^, representing a model of large vessel TES. In all experiments we also tested the enzymatically inactive Marburg I polymorphism (MI-SPD)^30^, which differs by a single amino acid, as a control. Both substances were infused together with tPA, the standard thrombolytic treatment, and compared to tPA alone and PBS (Fig. 1a). Since FSAP alone did not to exhibit any fibrinolytic activity^13^, WT-SPD was not tested on its own. Treatment infusions were started 60 min after the onset of thrombosis in the carotid artery. In pilot studies we observed a superior effect of 8.0, compared to 2.0, mg/kg body weight of FSAP-SPD and thus the higher concentration was used throughout the study. This is similar to the mode of delivery and amount of tPA used in such experiments^31^.

**Fig. 1:**
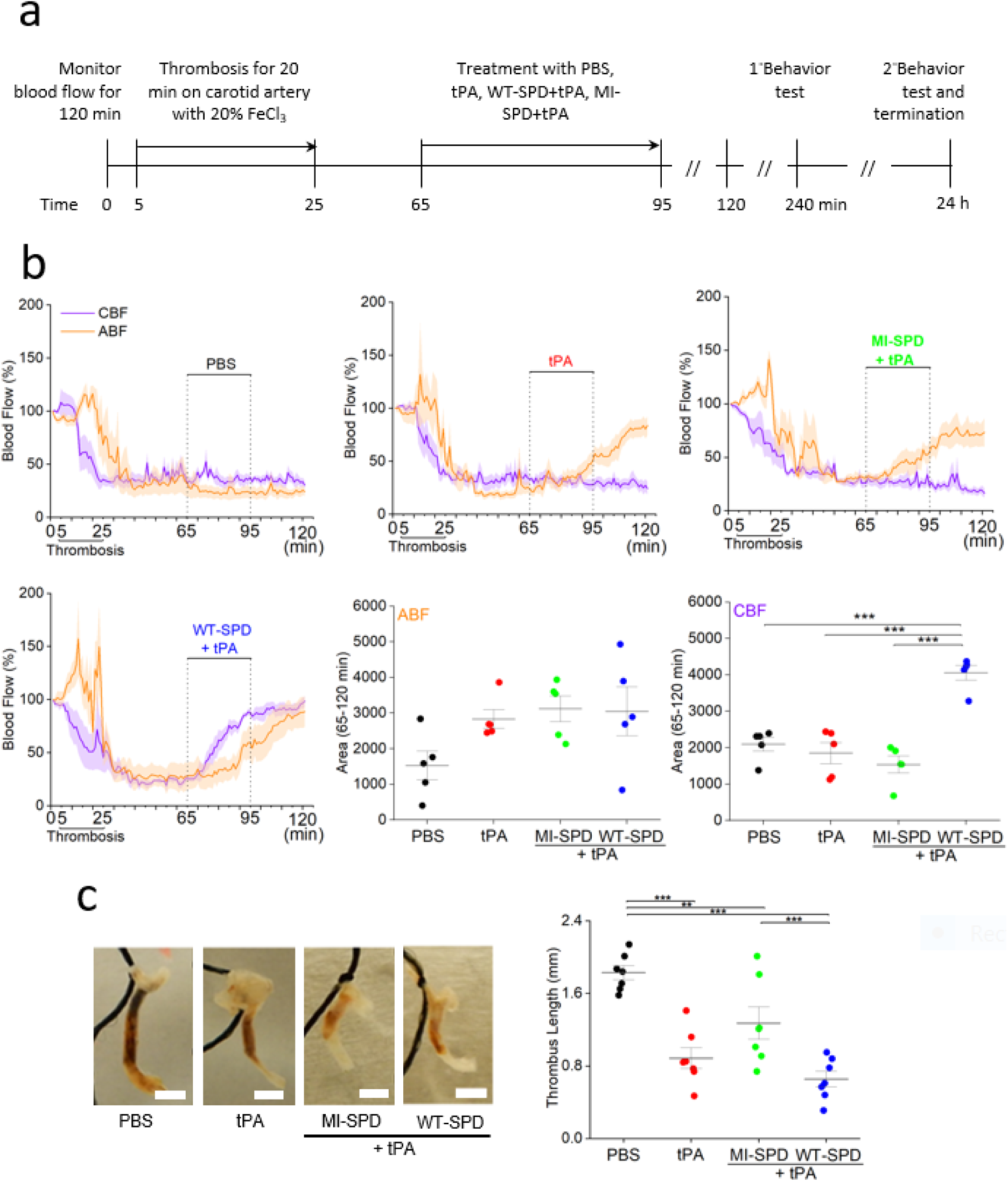
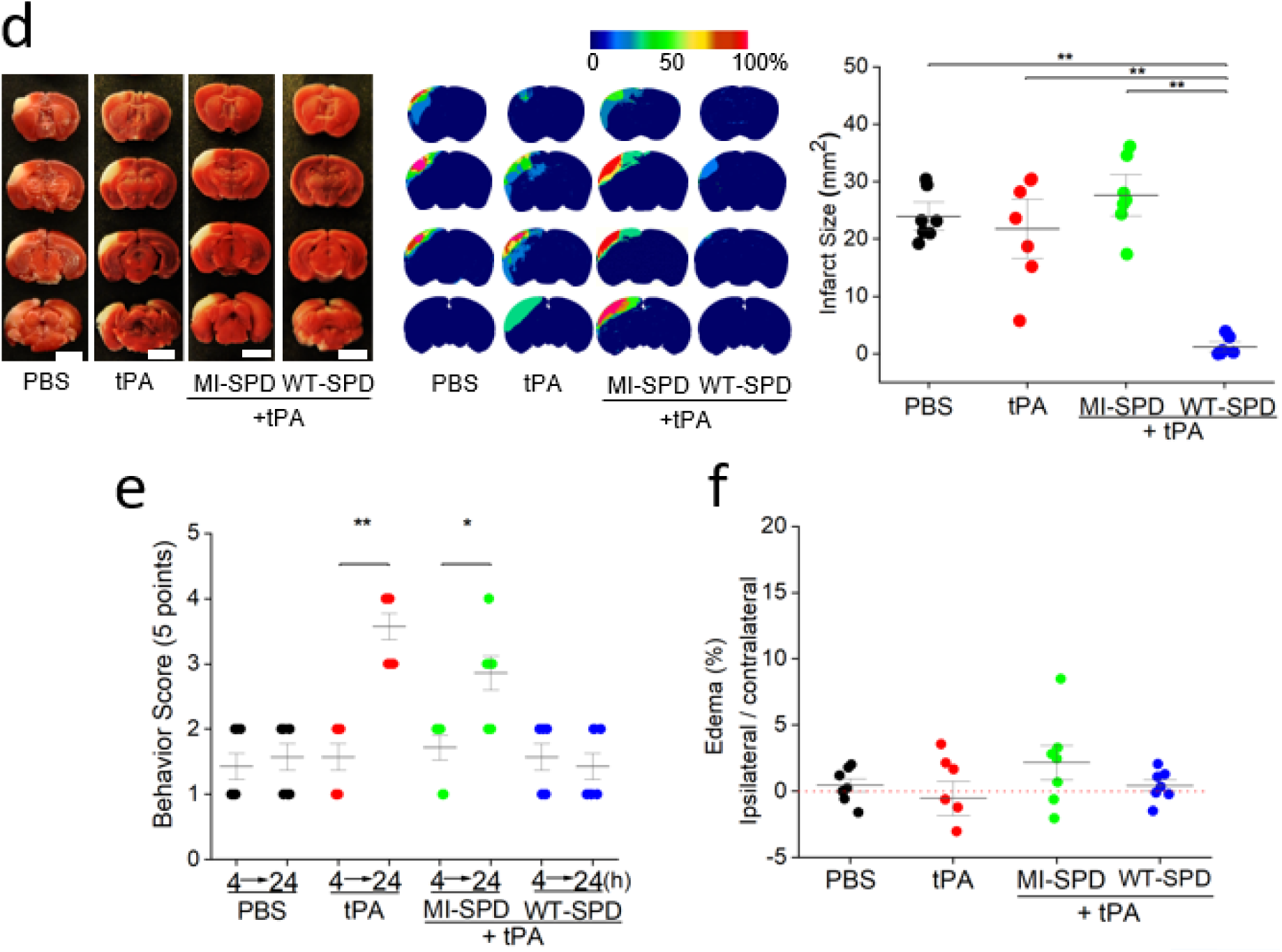
Effect of WT-SPD on the outcome of TES. **a**, Protocol of the experiment. **b**, Cerebral blood flow (CBF) and carotid artery blood flow (ABF) during the time course of the experiment after different treatments. Cumulative ABF and CBF between 65-120 min for each mouse is shown (n=5 mice per group). **c**, Images of carotid arteries from one mouse in each group and thrombus length (mm) (n=7 mice per group) (Scale bars = 1 mm). **d**, 2,3,5 triphenyltetrazolium chloride (TTC) staining of brain sections from one mouse in each group and pseudo colour representation of infarct size from all mice in the treatment group and its quantification (n=7 mice per group) (Scale bars, 5 mm). **e**. Behaviour score was measured with a 5-point scale (0-4) (n=7 mice per group). **f**, Edema-mediated change in size of ipsilateral over the contralateral hemisphere (n=7 mice per group). The data represents mean ± SEM and each dot represents one mouse. P-value was determined by one-way analysis of variance (ANOVA) with post-hoc Bonferroni test except in panel **e** where Kruskal-Wallis test with Mann-Whitney post-test was used; *p < 0.05, **p < 0.001, ***p < 0.0001.

We summarized that the initiation of thrombosis, and later thrombolysis, can be followed by measuring changes in perfusion in the carotid artery as well as the brain. Laser Doppler flowmetry was used to track the consequences of thrombosis and the subsequent fibrinolysis on cerebral blood flow (CBF) in the middle cerebral artery (MCA) territory as well as the carotid artery blood flow (ABF) downstream of the thrombus. The validity of this experimental design was first tested by temporarily ligating the carotid artery with a suture. This reduced CBF, whereas compensatory mechanisms increased ABF via collateral circulation as described before^32^ (Suppl. Fig, 1e). Thrombosis in the carotid artery reduced CBF, whereas ABF first increased transiently before decreasing (Fig. 1b). Treatment with tPA, tPA plus WT-SPD and tPA plus MI-SPD did not significantly influence CBF, but ABF was increased in all treatment arms with tPA (Fig 1b). However, only tPA plus WT-SPD increased CBF over time (Fig. 1b). We then assessed the size of the primary clot in the carotid artery. The size of the occluding clot was reduced in all groups with tPA treatment, confirming arterial thrombolysis, but the additional presence of either isoform of SPD did not influence the residual clot size significantly (Fig.1c). To consolidate these results, we also monitored blood flow in the neck and brain region using laser speckles contrast imaging (LSCI), and observed the same pattern of results with respect to ABF (Suppl. Fig. 2a,b) and CBF (Suppl. Fig. 2c,d). These changes in blood flow patterns need to be further correlated with the appearance and disappearance of clots in the brain.

The TES model mimics the often encountered clinical situation where tPA recanalizes the primary blocked artery, but fails to establish distal reperfusion^33^. The likely reasons for this resistance to thrombolysis are vascular dysfunction, cellular aggregation, biomechanical factors, ischemia-reperfusion injury and clot composition^34^ and FSAP might influence one or more of these pathways. Based on the known effects of FSAP on fibrinolysis^13^, we postulated that better clot lysis could account for the above results and studied the lysis of mouse blood clots *in vitro* to test this hypothesis. We found that the overall dissolution of clots with tPA plus WT-SPD was significantly better compared to using tPA alone or tPA plus MI-SPD (Suppl. Fig. 3). The time to maximal velocity of clot lysis was also faster in the presence of WT-SPD and tPA (Suppl. Fig. 3). WT- or MI-SPD alone, had no direct effect on clot lysis as reported earlier with human plasma-FSAP in human plasma clots^13^. Application of FSAP-SPD in mice lead to the formation of FSAP inhibitor complexes which were not investigated further (Suppl. Fig. 1A). Hence, an alternative explanation for faster lysis of whole blood clots in the presence of FSAP-SPD is that it scavenges fibrinolytic inhibitors such as plasminogen activator inhibitor-1^35^ and alpha2-antiplasmin^36^ and thus furthers the activity of tPA and plasmin.

We then analyzed the consequences of stroke after 24h in these mice. Similar infarct size, involving much of the dorsal and ventral striatum with overlying cortical areas, were observed in animal groups treated with PBS, tPA, and tPA with MI-SPD, while infarct size in mice treated with tPA plus WT-SPD were strikingly smaller (Fig. 1d). Behavioral scoring showed that mice treated with tPA performed worse after 24h, compared to the PBS, highlighting the adverse effects of tPA that are also observed in patients^37^. The scores were better in mice treated with tPA plus WT-SPD compared to tPA plus MI-SPD (Fig. 1e). There were no, edema-related, changes in hemisphere volumes (Fig. 1f). The mortality was 2/9 in PBS, 3/10 in tPA, 3/10 in tPA plus MI-SPD and 1/8 in tPA plus WT-SPD group (Fig. 1). These, non-random, attrition rates appear to be related to the treatments and can influence the effect size as has been described before^38^. Taken together, these findings show that better clot lysis with WT-SPD and tPA is associated with improved perfusion and reduced deleterious effects of tPA on neurological outcome and infarct volume. MI-SPD did not have any of these effects.

Considering the number of opposing actions FSAP has with respect to coagulation and fibrinolysis^8,12,13^, its application *in vivo* can lead to side effects related to these processes. We therefore examined the effects of FSAP-SPD on hemostasis in a tail-bleeding model (Fig. 2a). WT-SPD decreased the bleeding time (Fig. 2b) and the amount of blood loss (Fig. 2c) compared to MI-SPD or PBS treatment, which is in agreement with our earlier results^15^. Control mice treated with heparin bled profusely during the entire experiment (Fig. 2b,c). Since FSAP cleaves fibrinogen and promotes fibrinolysis, we also assessed the effect of the different treatments in the TES model on circulating levels of fibrinogen and d-dimer, which is a product of fibrinolysis (Fig. 2d)^39^. Fibrinogen levels were reduced in the tPA-treatment group compared to mice given PBS, but this reduction was not affected by the co-administration of WT-SPD (Fig. 2e). D-dimer exhibited the same trend (Fig 2f), which is a sign of lower coagulation and fibrinolysis and is usually associated with a better outcome in stroke^40^.

**Fig. 2:**
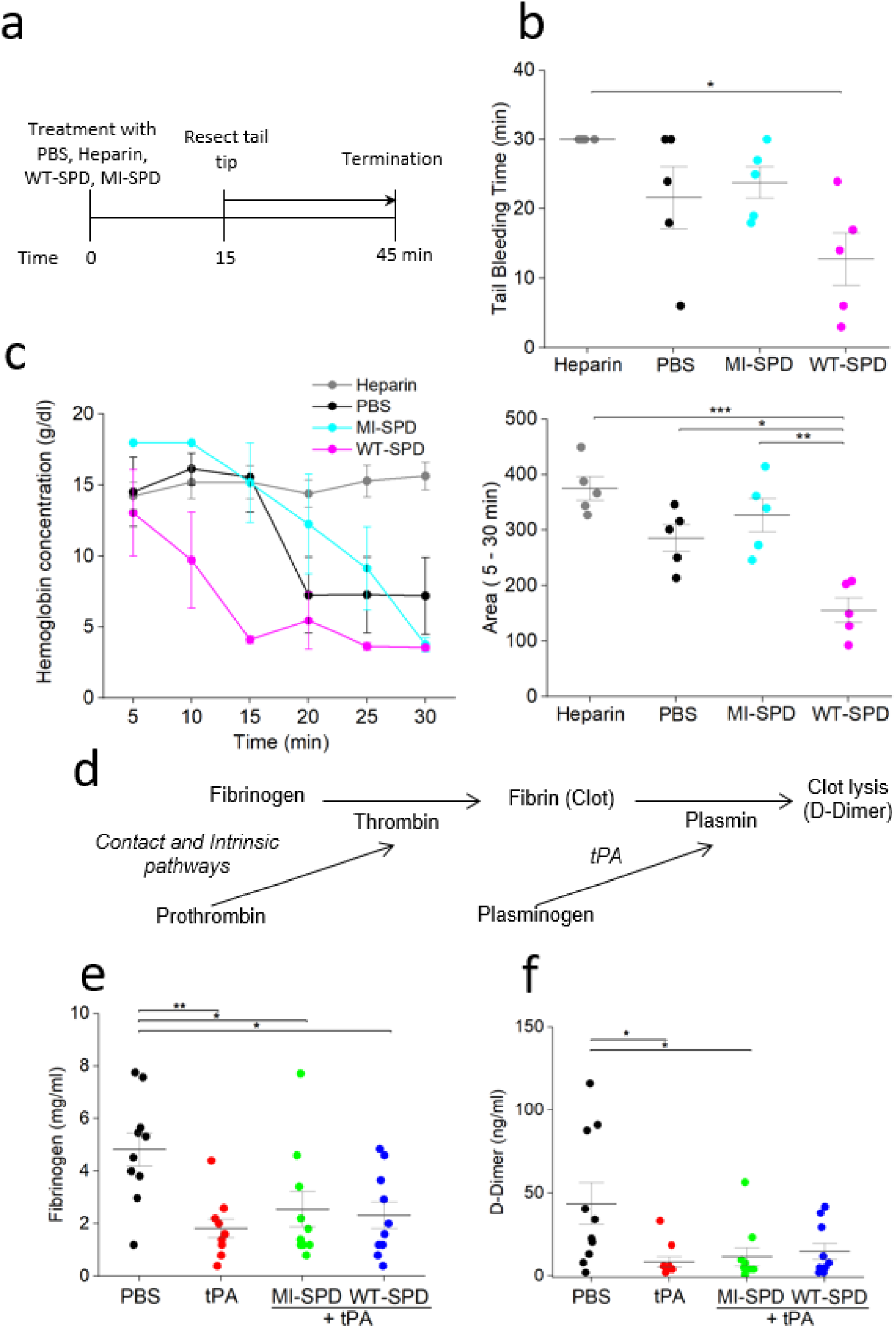
Effect of WT-SPD on hemostasis. **a**, Protocol of the experiment. **b, c**, Mice were treated with PBS, MI-SPD (8mg/ kg body weight), WT-SPD (8mg/ kg body weight) and heparin (10 IU/kg body weight) (n= 5 mice per group) and the time to stop bleeding from the resected tail was determined and blood loss was measured in 5 min intervals. Cumulative blood loss over the 30 min for each mouse was calculated **d**, Pathway for the formation of fibrin clot from fibrinogen and d-dimer after clot lysis **e, f**, Plasma fibrinogen (mg/ml) and d-dimer (ng/ml) was measured in citrate plasma 24-h after the induction of TES in the treatment groups described in Fig. 1a (n=10 mice per group). Each dot represents one mouse and the data represents mean ± SEM. P-value was determined by one-way analysis of variance (ANOVA) with post-hoc Bonferroni test; *p < 0.05, **p < 0.001, ***p < 0.0001.

WT-SPD provoked a hemostasis response, most probably, due to the inactivation of tissue factor pathway inhibitor^12,25^. The seemingly incongruent effects; the ability to promote coagulation and, at the same time, improve stroke outcome can be reconciled with the hypothesis that in the stroke model the pro-fibrinolytic effect of FSAP predominates over the pro-coagulant effect leading to an overall positive outcome. Through detailed structure-function studies it may be possible to mutate away such potentially negative effects as has been successfully done with other coagulation factors such as activated protein C (APC)^41^.

To test the generality of these finding we also tested the FSAP-SPD in the transient middle cerebral artery (MCA) occlusion (tMCAO) model of stroke. This model features ischemia-reperfusion injury and microvascular obstruction, rather than overt large vessel thrombosis^42^. In these mice, treatments were initiated 75 min after the onset of ischemia (Fig. 3a). CBF remained depressed in the reperfusion phase in all treatment groups (Suppl. Fig. 4a,b), most likely due to non-thrombotic obstruction of the microvessels. No differences in plasma-fibrinogen or d-dimer levels were observed (Suppl. Fig. 4c,d) confirming a lack of involvement of overt coagulation/ fibrinolysis in this model.

**Fig. 3:**
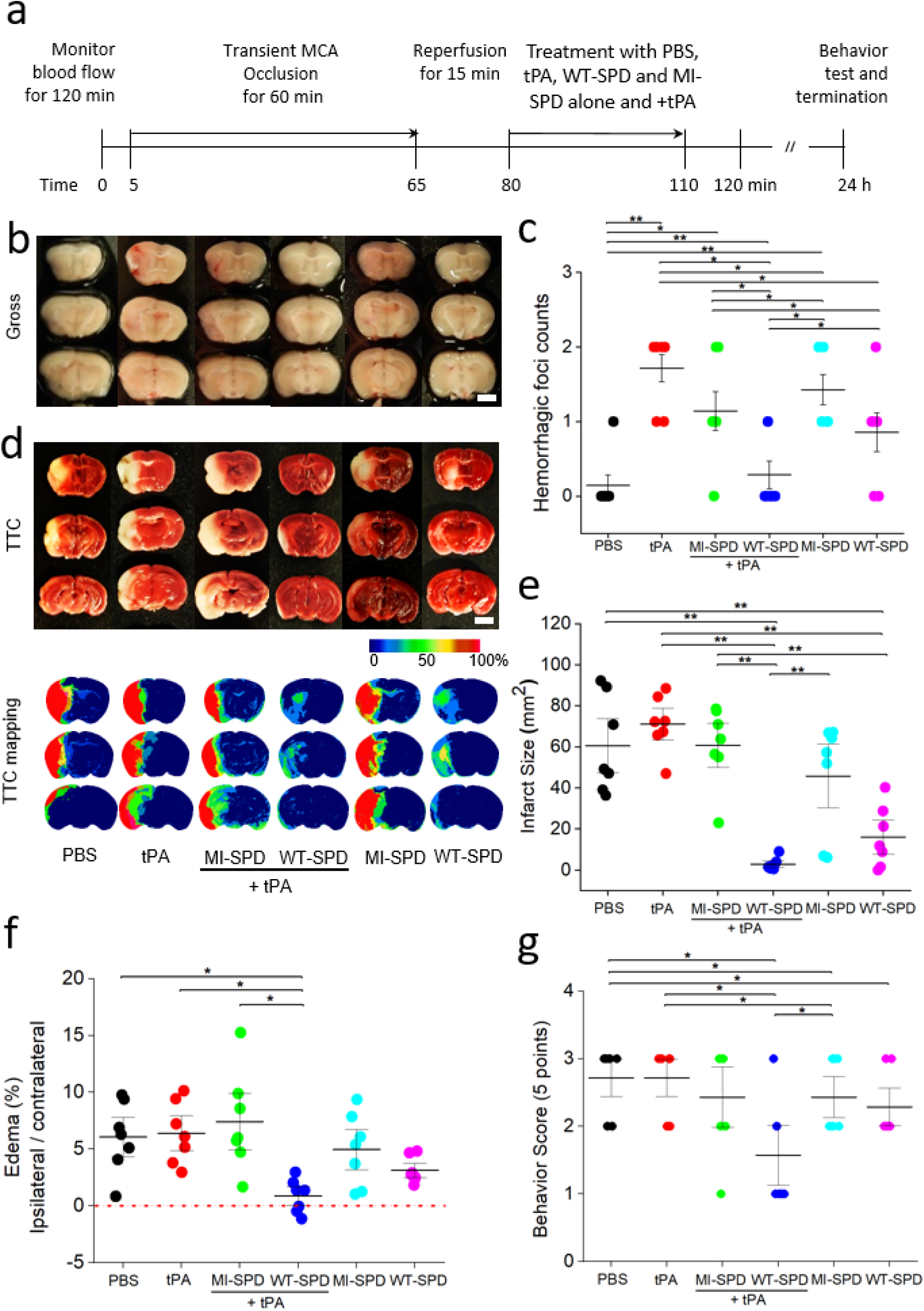
WT-SPD improves the outcome of stroke after tMCAO. **a**, Protocol of the experiment (n=7 mice per group in all panels). **b**,**c** Hemorrhagic foci from the brain sections from one mouse in each group and quantification. **d**,**e** TTC staining of brain sections from one mouse in each group and pseudo color representation of infarct size from all mice in the treatment group and quantification. **f**, The percentage change in the size of infarct hemisphere over the contralateral hemisphere due to edema. **g**, Neurological outcome of a 5-point scale (0-4). Each dot represents one mouse and data is mean ± SEM (n=7). P-value was determined by one-way analysis of variance (ANOVA) with post-hoc Bonferroni test except in panel **c** and **g** where Kruskal-Wallis test with Mann-Whitney post-test was used; *p < 0.05, **p < 0.001, ***p < 0.0001. In **b** and **d**, scale bar, 5 mm.

We then analyzed the consequences of stroke after 24h in these mice. We found that all stroke parameters were significantly improved in the WT-SPD plus tPA groups, compared to the other groups. The number of hemorrhagic foci were increased in tPA-treated mice, while only to a lesser extent in mice also receiving WT-SPD. For unknown reasons the number of foci were significantly elevated in the MI-SPD group (Fig 3b,c). The infarct size was also larger in groups treated with tPA, MI-SPD, or their combination, while being significantly smaller in mice that received WT-SPD alone or together with tPA (Fig 3d,e). There was a substantial increase in the edema-related expansion of the stroke-hemisphere in all mice, except in the WT-SPD plus tPA treated groups (Fig. 3f), as were the neurological deficit scores (Fig. 3g). The mortality was 2/9 in PBS, 2/9 in tPA, 2/9 in tPA plus MI-SPD, 2/9 in tPA plus WT-SPD, 3/10 in MI-SPD and 1/8 in WT-SPD group (Fig. 3a-g). Hence, in a model with a minor involvement of coagulation pathway, FSAP-SPD had a beneficial effect on its own as well as reduced the deleterious effects of tPA.

In earlier studies, FSAP deficiency (*Habp2*^-/-^) worsened the outcome of experimental stroke^11^ in a model that relied on a direct injection of thrombin into the MCA to induce thrombosis. The role of endogenous FSAP has not been systematically tested in other models of ischemic stroke. In the tMCAO model these mice exhibited a small, but significant, increase in infarct volume, but there were no differences in the TES model (Fig. 4a-d). Treatment of *Habp2*^-/-^ mice with FSAP-SPD in the TES model led to a similar outcome as observed in WT mice (Fig. 4e-h). In general the exclusion rates were higher in experiments with *Habp2*^-/-^ mice compared to WT mice. One explanation for this could be that thrombi in these mice have a higher propensity to embolize, as described before^15^, and disintegrate more easily upon handling (Fig. 4C) which could influence the survival of the mice in stroke models.

**Fig. 4:**
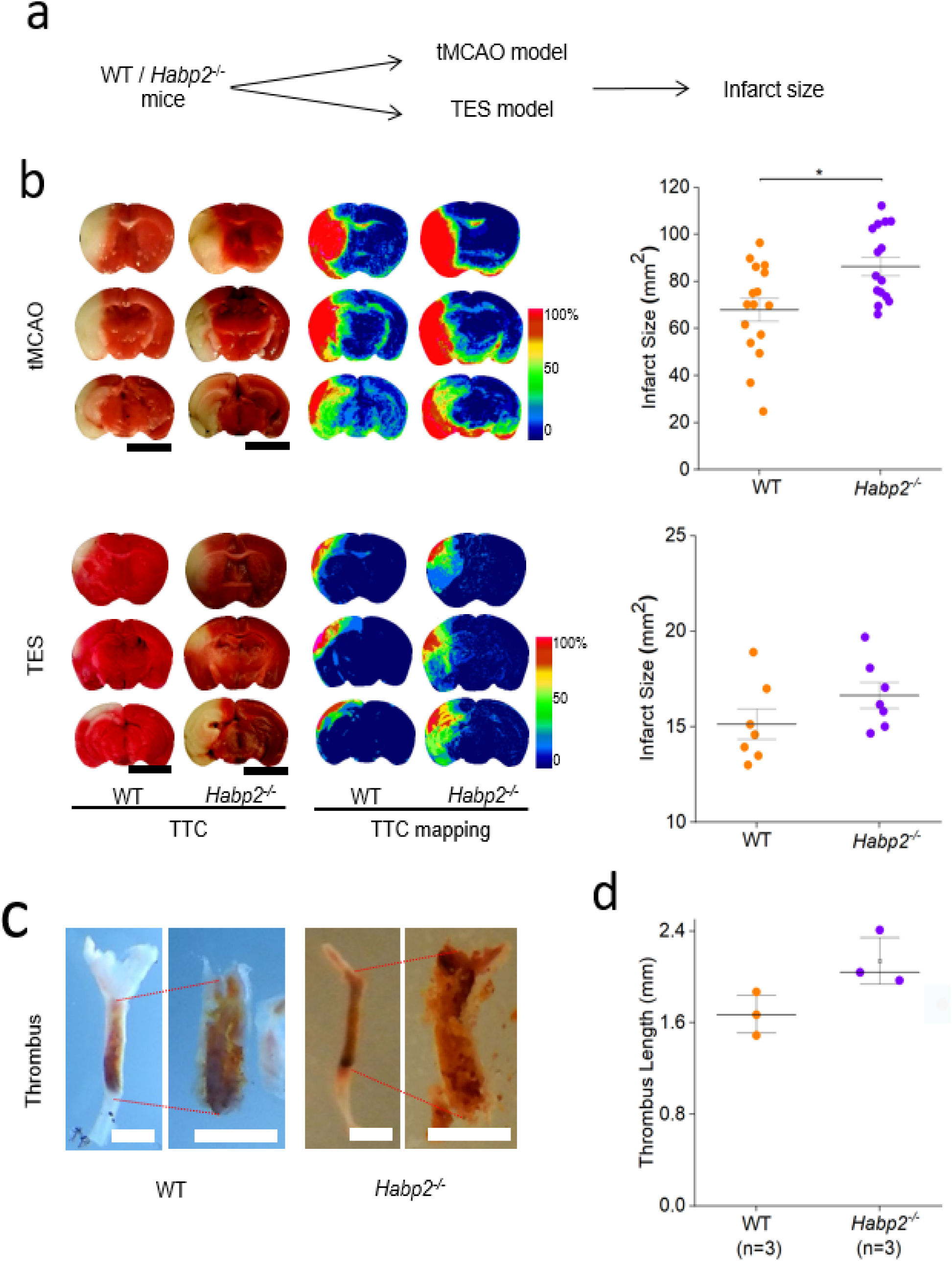
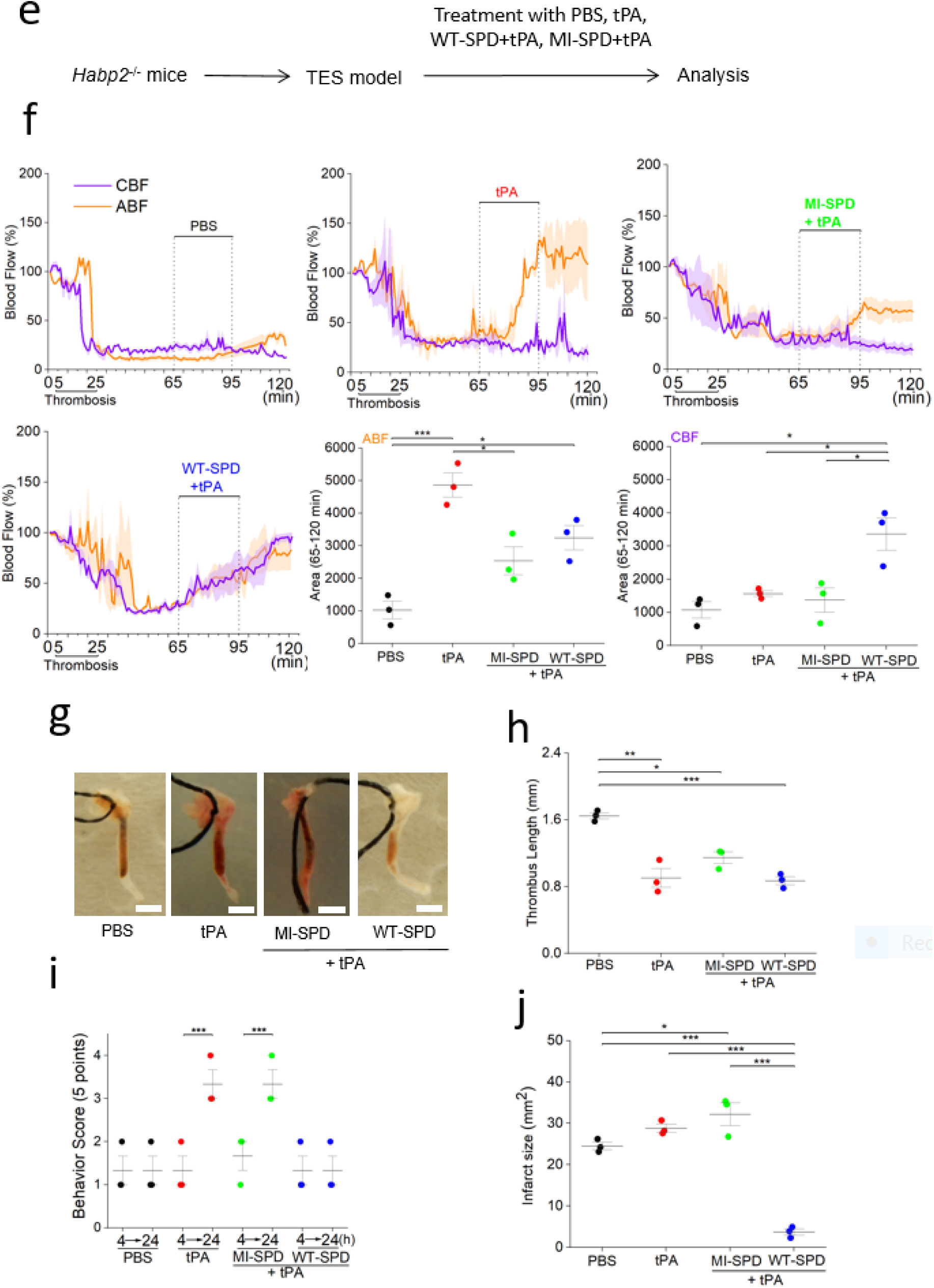
Outcome of TES and tMCAO in *Habp2*-/- mice. **a**, Outline of the experiment. **b**, Images of consecutive TTC-stained sections of WT and *Habp2*^-/-^ mice in both model of stroke, pseudo color mapping of sections from all mice within the group and quantification of infarct size (scale bar = 5 mm). Each dot represents one mouse (n=16 mice in tMCAO model and n=7 mice in TES model). Results from two independent experiments are pooled in the tMCAO panel. **c**,**d**, Images of carotid arteries from one mouse in each group and thrombus length (mm)(n=3 mice per group) (scale bars =1 mm). **e**, *Habp2*^-/-^ mice were subjected to TES stroke model and treated as described in Fig. 1a. **f**, Percentage of basal CBF and ABF during the experiment after treatment with PBS, tPA, tPA plus MI-SPD or tPA plus WT-SPD. Cumulative ABF and CBF between 65-120 min for each mouse (n=3 mice per group). **g**, Images of carotid arteries from one mouse in each group and **h**, thrombus length (mm) (n=3 mice per group) (scale bars = 1 mm). **i**, Behaviour score was measured with a 5-point scale (0-5) (n=3 mice per group). **j**, The infarct size from TTC-stained sections (n=3 mice per group). Each dot represents one mouse and the data represent mean ± SEM. P-value was determined by one-way analysis of variance (ANOVA) with post-hoc Bonferroni test except in panel **i** where Kruskal-Wallis test with Mann-Whitney post-test was used; *p < 0.05, **p < 0.001, ***p < 0.0001.

## Conclusions

Pro-thrombolytic drugs typically have problems^13^ related to increased hemorrhage and/or bleeding associated with increased tPA and/or plasmin activity^37^. An advantage of using FSAP is that it promotes fibrinolysis without altering tPA or plasmin activity since it acts directly on fibrin/ogen. On the other hand, the pro-coagulant effects of FSAP, observed here and in a previous study^15^, are possibly problematic and theoretically could worsen the outcome of stroke. However, this was not observed in both the stroke models indicating that this may be a weak side effect of FSAP. Counterintuitively, this pro-coagulant effect might negate the hemorrhagic transformation due to tPA. FSAP may also directly inhibit the known negative effects of tPA treatment^43^ independently of its effects on hemostasis.

A recent study by Tian et al showed that high molecular weight hyaluronic acid (HMW-HA) down-regulated FSAP *in vivo* and in endothelial cells *in vitro* and improved the outcome of stroke in the tMCAO model^9^. The interpretation made is that the presence of excessive FSAP is detrimental for stroke outcome, which is the opposite of our conclusion. However, it should be noted that HMW-HA treatment causes extensive changes in the gene expression profile of endothelial cells^9^ and in lieu of this fact it is not possible to ascribe the positive effects of HMW-HA treatment on endothelial dysfunction exclusively to a reduction in FSAP alone.

Limitations of the current study are that we used the serine protease domain of FSAP and not the full length protein. The full-length protein maintains the full complement of regulatory interactions which are lost with the use of the SPD domain thus limiting the extrapolation of these results to a physiological context. The use of human protein in the mouse may lead to aberrant conclusions that need to be replicated by using recombinant mouse FSAP. However, the same expression system, used for producing human FSAP-SPD, was not suitable for mouse FSAP-SPD expression (data not shown). The effects of FSAP were not tested in aged or female mice, or in mice with co-morbidities which are important factors in determining drug efficacy in stroke treatment as described in the STAIRS guidelines^44^. The study did not consider the long term effects of FSAP-SPD treatment or FSAP deficiency. Molecular mechanisms of action with respect to PAR activation, apoptosis, vascular permeability and histone detoxification were not further elucidated *in vivo* in more detail.

This study is the first reported use of recombinant FSAP-SPD as a direct therapeutic agent in ischemic stroke in animal models. These results are congruent with the observations in relation to FSAP and stroke in humans and in FSAP-deficient mice. FSAP has the advantage of being an endogenous blood protein with a plausible mechanism of action that is readily amenable to further development.

## METHODS

### Mice

Adult male C57BL6J mice (10-15 weeks old) were obtained from Charles Rivers (Sulzfeld, Germany) and FSAP-deficient mice (*Habp2*^-/-^, B6129S5-*Habp2*^tm1Lex^)^45^ on a C57BL6J background were bred in our institution. All mice were kept under specific pathogen-free (SPF) conditions. All animal experiments were approved by the governing body regulating research on animals in Norway (Forsøksdyrforvaltningens tilsynsog søknadssystem (FOTS); Application Nos. 7557, 9769 and 9772). Group size was calculated for the effect of treatment on infarct area 24 h after stroke in WT mice. We assumed a 40% difference in mean, 30% standard deviation, 80% power and 95% confidence which gave a group size of 7. We expected a premature termination of experiment in about 20-25% of the animals due to death before 24h or euthanasia due to excessive pain or moribund state. Because there were differences in attrition rates between the groups we operated 8-10 animals per group. Group size was increased for *Habp2*^*-/-*^ mice due to a higher attrition (40-50%) in this strain. Group size was not calculated for experiments related to blood flow, hemostasis or pharmacokinetics.

### TES model

Studies were performed in accordance with the “ARRIVE Essential 10” guidelines^46^ except that the operator was not blinded to the treatments. Mice were randomly assigned to different treatment groups (PBS, tPA, WT-SPD plus tPA or MI-SPD plus tPA). Mice were anesthetized with isoflurane and the core temperature was maintained at 37°C with a rectal-probe temperature-controlled homeothermic blanket (Panlab, Barcelona, Spain). Cerebral blood flow (CBF) was measured by a laser Doppler flowmeter (Moor, Axminster, UK) with a flexible fiber-optic probe affixed to the skull over the parietal cortex perfused by the MCA after making an incision in the skin. The neck area was dissected to expose the carotid artery and another fiber-optic probe was attached under the carotid artery bifurcation to measure arterial blood flow (ABF). After exposing the carotid artery a 1 × 1 mm^2^ filter paper soaked in 20% FeCl3 (w/v) solution was placed on the central section of the carotid artery for 20 min to induce thrombosis and stroke as described before^29^. The treatment with tPA (Altepase, Boehringer Ingelheim, Ingelheim, Germany), (10 mg/kg body weight) and SPDs (8 mg/kg/body weight), combinations of tPA and SPD as well as PBS was initiated as an intravenous infusion over 30 min (10% bolus) through the dorsal penial vein. The neck and head wound was sutured and the animals were transferred to temperature-controlled recovery cages and at 4 and 24h a 5-point behavior score was determined^47^. Normal motor function was scored as 0, flexion of the contralateral torso and forelimb on lifting the animal by the tail as 1, circling to the contralateral side but normal posture at rest as 2, leaning to the contralateral side at rest as 3, and no spontaneous motor activity as 4.

### tMCAO model

Mice were randomly assigned to different treatment groups; PBS, tPA, WT-SPD plus tPA, MI-SPD plus tPA, WT-SPD or MI-SPD. All procedures were the same as in the TES model except that after the neck area was dissected to free the carotid artery, a filament (0.22 ∼ 0.24 mm, 60-2356 PK10, Doccol corporation, Sharon, MA) was inserted through an incision in the external carotid artery and advanced through the internal carotid artery to the origin of MCA as described before^48^. After confirming occlusion by a 70% drop in CBF, the filament was kept in position for 60 min while the mice were maintained under anesthesia. After a 15 min reperfusion period the treatment was initiated and the rest of the procedures were as described under the TES model.

### Fibrinogen and D-dimer ELISA

Citrate plasma was collected from mice from the TES and tMCAO experiments before sacrifice at the 24h time point. Asserchrome D-dimer ELISA (#100947) from Stago (Asnières sur Seine, France) and Mouse total fibrinogen (#IMSFFBGKTT) from Innovative Research (Novi, MI) was used to analyse the samples in duplicates.

### Analysis of hemorrhage, infarct size, and hemisphere swelling

Animals were sacrificed at the 24h time point, the brains were retrieved and cut in 2 mm coronal sections on a brain matrix and photographed. Infarct size was determined from sections stained with 2,3,5 triphenyltetrazolium chloride (TTC) (2% w/v) in PBS at 37°C for 30 min. The areas of both the hemispheres and the infarct area (white) was measured using Photoshop and the infarct size was calculated. TTC staining was pseudo-colored to combine all images from all mice within a group using Image J. Foci of hemorrhage were counted on one face of the brain slices. The swelling of the infarcted hemisphere was determined by the formula: ((entire brain-infarct hemisphere × 100) / infarct hemisphere - contralateral hemisphere)).

### Laser Speckles contrast imaging (LSCI)

These experiments were performed on a separate set of mice with the same treatment groups as described above (n=3 mice per group in TES model and n=5 mice per group in tMCAO model). Blood flow was imaged over the whole neck region and the brain using a laser speckles imager (Moor FLPI-2, Axminster, UK). To image the brain, a midline incision was made on the scalp, and pericranium was removed to expose the surface of the skull in anesthetized mice. Images were aquired under laser illumination at the indicated time points and were pseudo-colored and blood flow is expressed as percentage of baseline.

### Tail resection model

The effect of WT-SPD, MI-SPD, Heparin and PBS on hemostasis was tested. Isoflurane-anesthetized mice (n=5 mice per group) were given different treatments i.v. through the dorsal penile vein and 15 min later 0.5 cm tip of the tail was resected before the tail was placed in a PBS at 37°C. The PBS solution was changed every 5 min for 30 min and the time to stop bleeding was noted. The experiment was terminated after 30 min by sacrificing the animals. Hemoglobin concentration in the PBS solutions was measured using the Hemoglobin Assay kit (Merck, MAK115).

### In vitro clot lysis experiments

Whole blood from C57BL6J mice (n=9 mice) without anticoagulants was directly collected into a polyethylene tube (PE-50, I.D. 0.58 mm) and allowed to clot for 2 h at room temperature. Thereafter the tubing was cut into 2 cm sections and the clots ejected into wells of a microtiter plate with PBS (triplicate wells per treatment). Clot lysis was initiated with tPA (20 μg/ml) in 10% plasma from other donor mice in the absence of presence of WT-SPD or MI-SPD (10 μg/ml) and the liberation of erythrocytes/ hemoglobin from the clots was measured at 37°C at 510 nm in a plate reader (Biotek). Time to maximal velocity of clot lysis and the end absorbance were quantified across different experiments.

### FSAP-SPD preparation and pharmacokinetics

The C-terminal His tagged serine protease domain (SPD) of human WT-FSAP and the MI mutation was prepared and characterized as described before^25^. Mice (n=3 mice) were given WT-SPD at 12 mg/kg body weight intravenously as a bolus and citrate-plasma (0.38% w/v citrate) was collected at various time points afterwards. Ac-Pro-*D*Tyr-Lys-Arg-AMC) (amino-methyl-coumarin) was used as a sensitive and specific substrate for human FSAP as described before^26^. Hydrolysis of the fluorogenic substrates was measured using a plate reader with excitation at 320 nm and emission at 460 nm (37°C for 60 min) at 37°C. The maximal velocity was calculated from the linear part of the progress curve. An enzyme-linked immunoassay (ELISA) was used to determine the concentration of FSAP-SPD and Western blotting with anti-FSAP antibodies was also used to monitor plasma levels^26^. Antibody for Western blotting and ELISA is specific for human FSAP with no or little cross reactivity with mouse FSAP.

### Statistical analysis

Data are expressed as mean ± SEM. Results from each individual mouse are shown as a point in all panels. Data were analysed by one-way analysis of variance (ANOVA) with post-hoc Bonferroni test for continuous measures; *p < 0.05 and **p < 0.001 ***p < 0.0001. For categorical measures we used Kruskal-Walis test with Mann-Whitney post-test. SSPS statistical package was used for all analysis (IBM, SPSS 26).

## Acknowledgements

Research funding was from the Research Council of Norway [Grant # 251239] to SMK and The Scientia Fellows programme of the University of Oslo in conjunction with the Marie Sklodowska Curie programme of Horizon 2020 to JYK. Additional funding was provided by the European Union’s Horizon 2020 Framework Programme for Research and Innovation under the Specific Grant Agreement No. 785907 (Human Brain Project SGA2), Specific Grant Agreement No. 945539 (Human Brain Project SGA3), and The Research Council of Norway under Grant Agreement No. 269774 (INCF Norwegian Node) to TBL.

## Author contributions

JYK performed all of the animal experiments, the imaging studies, analysed the data, performed the statistics and prepared the figures. DM prepared and characterized the recombinant proteins. TBL supervised the morphological analyses. SMK designed the study, obtained funding, analysed the data and drafted the manuscript. All authors read, edited and approved the final version of the manuscript.

## Supplementary figures and legends

**Supplementary Fig. 1:**
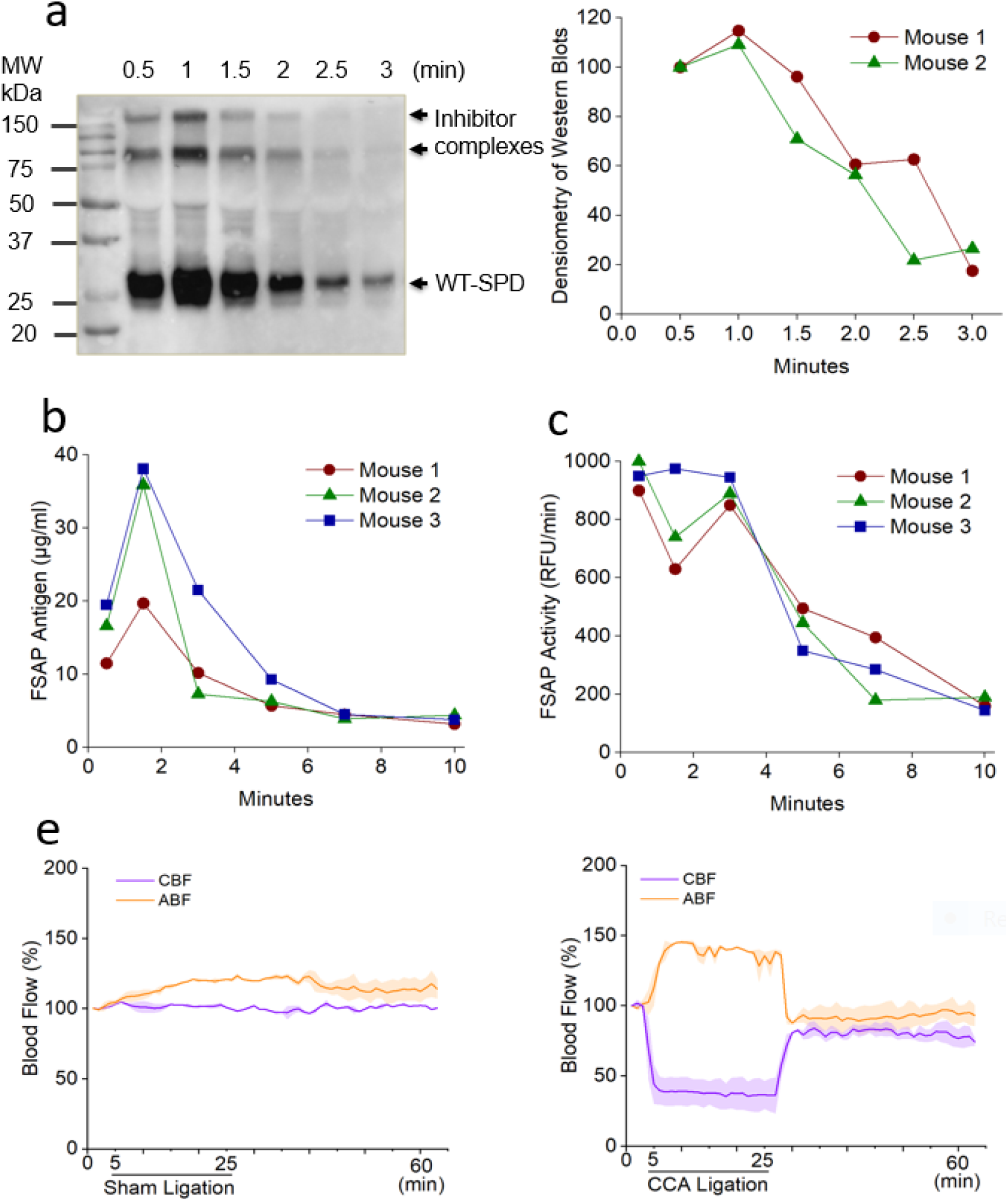
Pharmacokinetics of WT-SPD in mice *in vivo* and blood flow changes in the artery and brain after temporary ligation of the common carotid artery (CCA). **a-d**, WT-SPD (12 mg/kg body weight) was injected in the tail vein and plasma was collected at different time points (n=3 mice per group). **a**, FSAP antigen was detected by Western blotting with a rabbit anti-FSAP antibody (n=2 mice per group) and densiometric quantification. **b**, FSAP ELISA or **c**, activity assays with Ac-Ala-Lys-Nle-Arg-AMC fluorogenic substrate in duplicate wells (n=3 mice per group). **d**, Cerebral blood flow (CBF) and carotid artery blood flow (ABF) during the time course of sham operation (left) and temporary CCA ligation with a suture (right) (n=3 mice per group).

**Supplementary Fig. 2:**
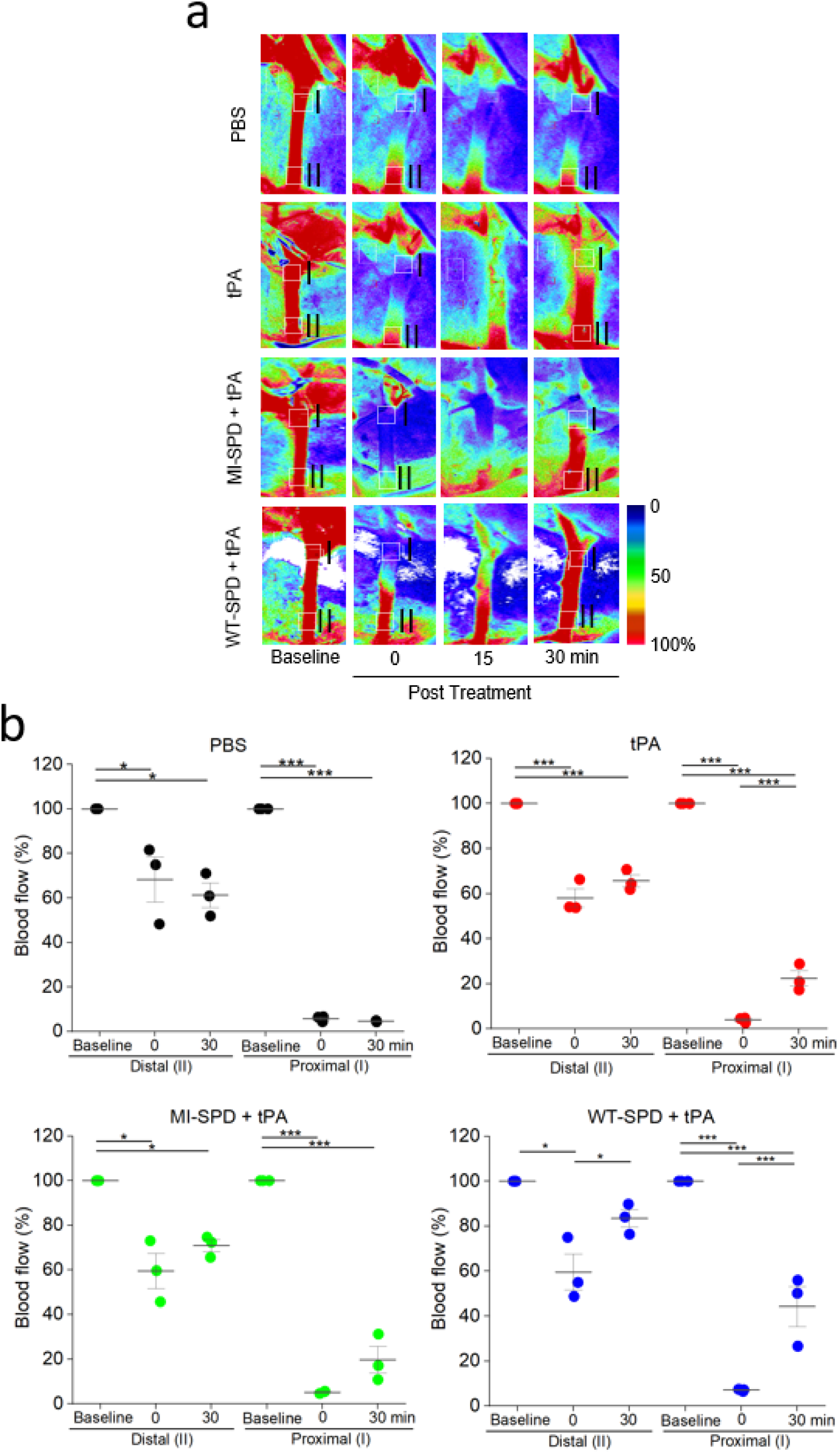
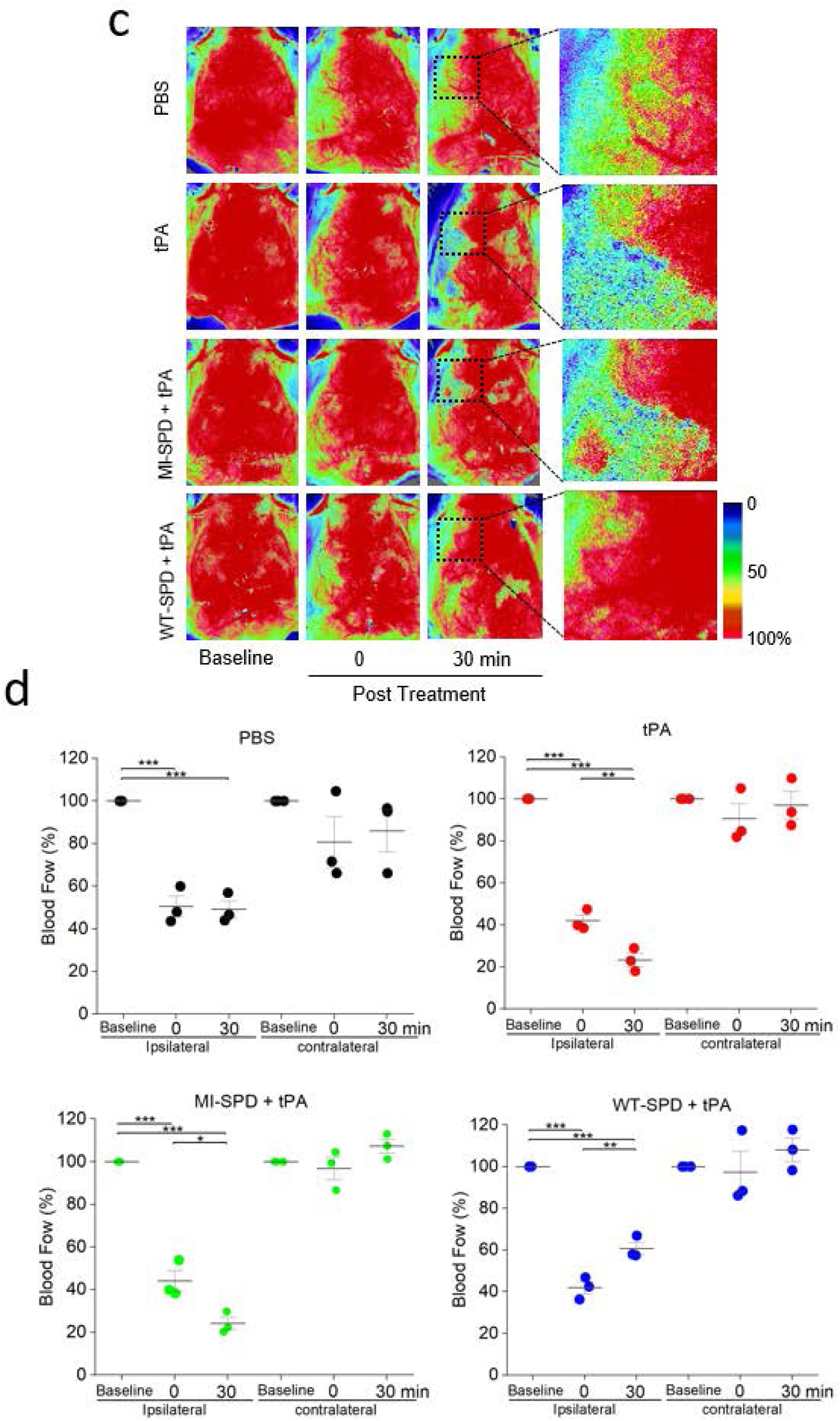
Arterial and cerebral blood flow after thromboembolic stroke (TES) as determined by laser speckles contrast imaging (LSCI). Protocol outline as in Fig. 1a of the manuscript. **a**, Carotid artery blood flow was measured during the time course of the experiment by LSCI after treatment with PBS, tPA, tPA plus MI-SPD or tPA plus WT-SPD. Images from one mouse in each group at baselines, 0, 15 and 30 min after initiation of treatment. Colour bar represents blood flow in arbitrary units. **b**, Quantification of the LSCI signal in the region of interest in the artery distal to the thrombus near the bifurcation of the carotid artery (I) and proximal to the thrombus (II). **c**, LSCI images from one mouse were recorded from the whole brain at 0 and 30 min after the initiation of treatment. **d**, Blood flow in the region of interest in the ipsilateral and corresponding region of the contralateral hemisphere was quantified. In **b** and **d** each dot represents one mouse and the data represent mean ± SEM (n= 3 mice per group). P-value was determined by one-way analysis of variance (ANOVA) with post-hoc Bonferroni test; *p < 0.05, **p < 0.001, ***p < 0.0001.

**Supplementary Fig. 3:**
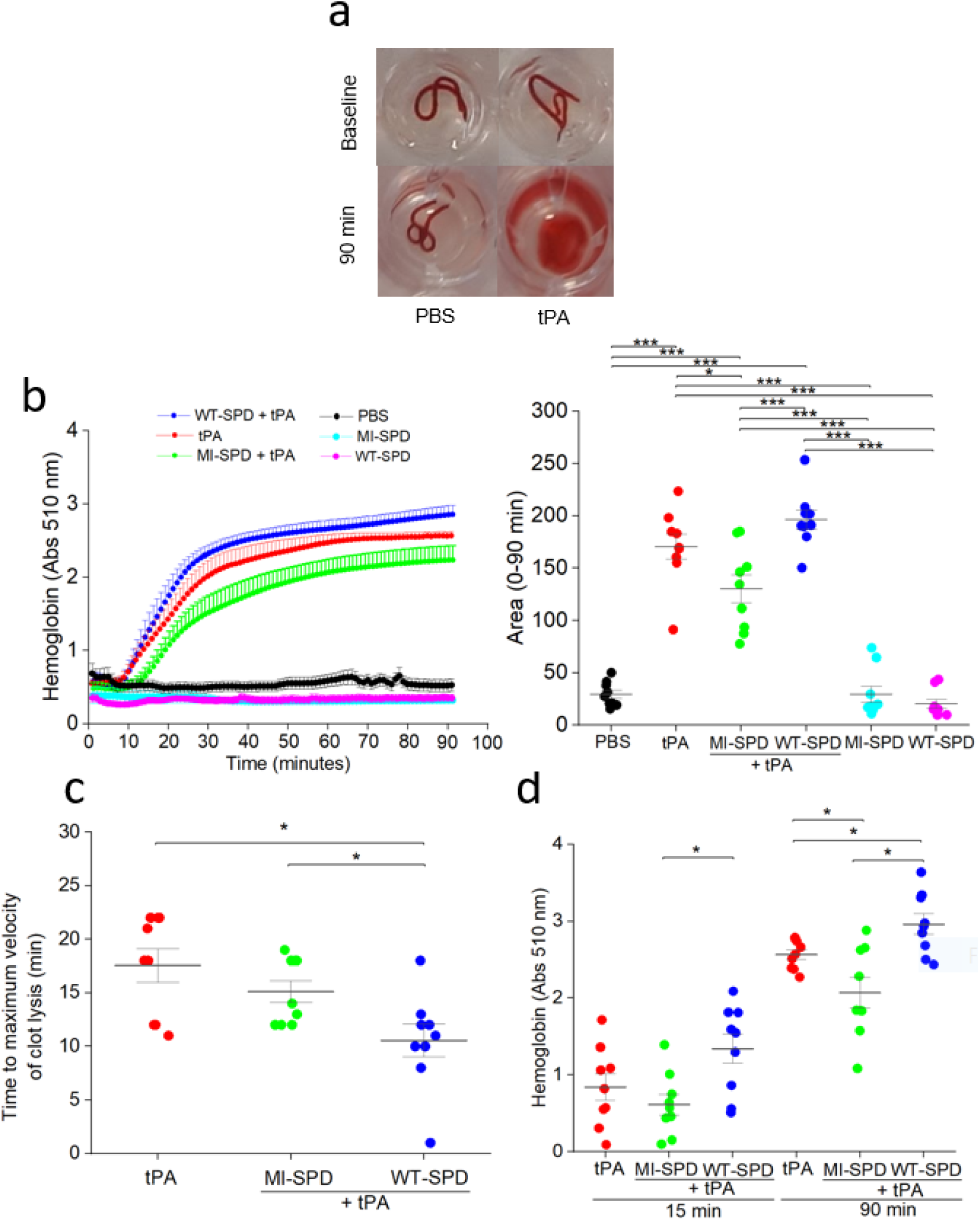
Mouse whole blood clot lysis by tPA in vitro. Whole blood clots (n=9 mice) formed *in vitro* were treated with PBS, tPA (20 μg/ml), WT-SPD (10 μg/ml) or MI-SPD (10 μg/ml) or their combinations for 90 min in the presence of 10% mouse plasma. **a**, Images of single wells at the start and the end of the experiment. **b**, Absorbance at 510 nm over 90 min in the different treatment groups from triplicate wells and area under the curve from 0-90 min in each experiment. **c**, Time to reach maximal velocity for clot lysis. **d**, Final absorbance at 90 min was measured to determine the total extent of clot lysis. Each dot represents clot lysis experiment on clots from 1 mouse and the data represent mean ± SEM (n= 9). P-value was determined by one-way analysis of variance (ANOVA) with post-hoc Bonferroni test; *p < 0.05, **p < 0.001, ***p < 0.0001.

**Supplementary Fig. 4:**
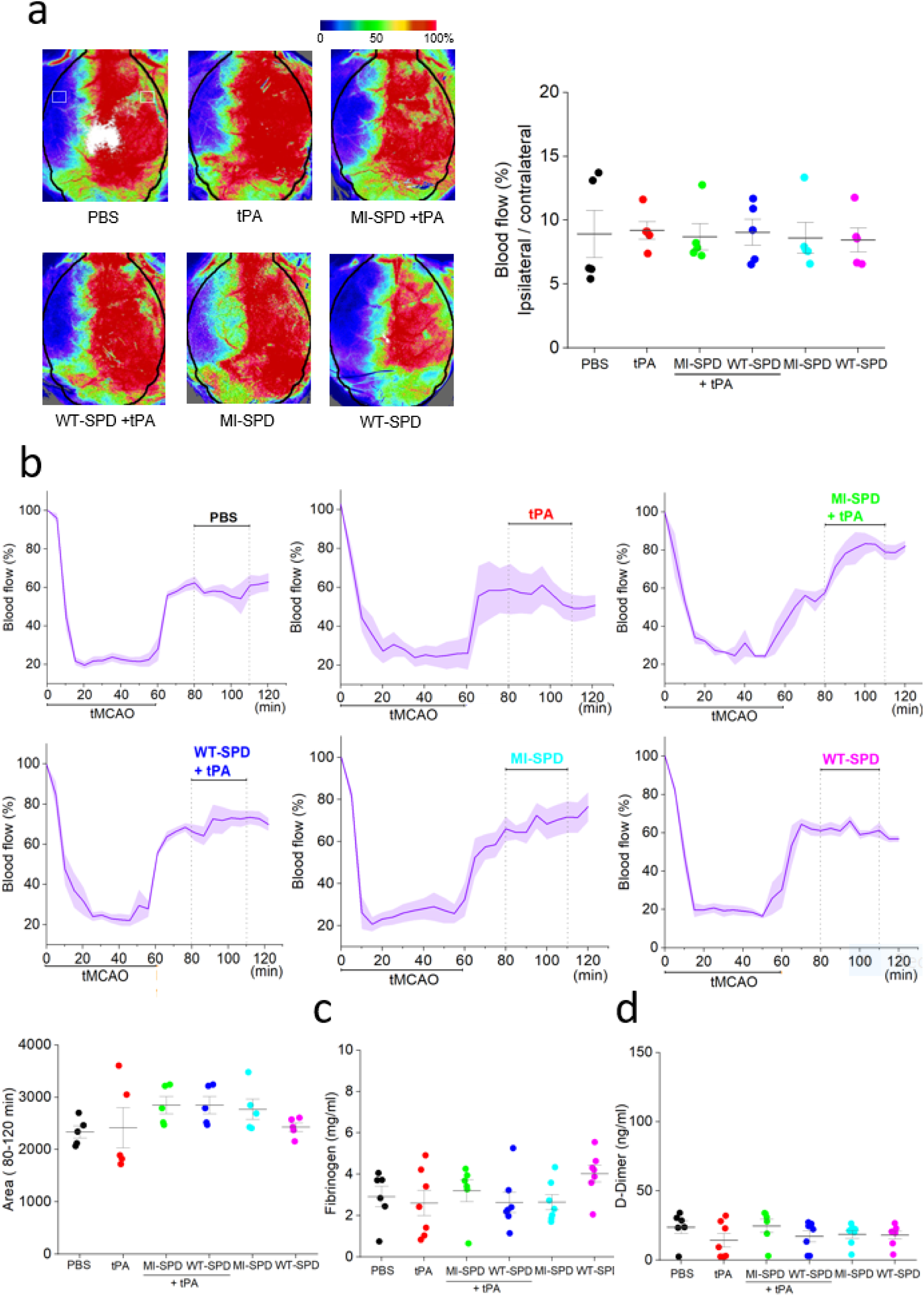
Brain blood flow changes during the tMCAO procedure using LSCI and laser Doppler flowmetry. **a**, CBF was measured by LSCI 60 min after MCA occlusion and image from one mouse per group is shown and expressed as percentage in the boxed region of ipsilateral / contralateral hemisphere (n=5 mice per group). **b**, CBF was measured using laser Doppler flowmetry over the 120 min period. Cumulative CBF in individual mice (bottom row, left panel) after the start of treatment (80-120 min) (n=5 mice per group). **c**,**d** Plasma fibrinogen (mg/ml) and d-dimer (ng/ml) was measured in citrate plasma 24-h after the induction of stroke in the treatment groups described in Fig. 3a (n=10 mice per group). Each dot represents one mouse and the data represent mean ± SEM. P-value was determined by one-way analysis of variance (ANOVA) with post-hoc Bonferroni test; *p < 0.05, **p < 0.001, ***p < 0.0001.

